# NRX-101, A RAPID-ACTING ANTI-DEPRESSANT, DOES NOT CAUSE NEUROTOXICITY FOLLOWING KETAMINE ADMINISTRATION IN PRECLINICAL MODELS

**DOI:** 10.1101/2022.06.18.496662

**Authors:** William Jordan, Richard Siegel, Rajendra Kumar, Jonathan Javitt

## Abstract

**Background:** NMDA (N-Methyl-D aspartate) receptor antagonists have gained increasing attention as rapid-acting antidepressants. However, their use has been limited by potential neurotoxicity (Olney Lesions) and recent FDA guidance requires demonstration of safety on histologic parameters prior to initiation of human studies. D-cycloserine is a mixed NMDA agonist/antagonist awarded Breakthrough Therapy Designation and currently in clinical trials for the treatment of bipolar depression with suicidal ideation. The current study was designed to investigate the neurologic safety profile of D-cycloserine by itself and in combination with ketamine and lurasidone.

**Methods:** Sprague Dawley female rats (n=106) were randomly divided into 7 study groups. Ketamine was administered via tail vein infusion. D-cycloserine and lurasidone were administered via oral gavage in escalating doses to a maximum of 2000 mg/kg DCS. To ascertain toxicity, dose escalation with three different doses of D-cycloserine/lurasidone was given in combination with ketamine. MK-801, a known neurotoxic NMDA antagonist, was administered as a positive control. Brain tissue was sectioned and stained with H&E and Fluorojade stains.

**Results:** No fatalities were observed in any group. No microscopic abnormalities were found in the brain of animal subjects given ketamine, ketamine followed by DCS/lurasidone, or DCS/lurasidone alone. Neuronal necrosis, as expected, was seen in the MK-801 positive control group.

**Conclusion:** NRX-101, a fixed-dose combination of D-cycloserine/lurasidone, when administered with or without prior infusion of IV ketamine was tolerated and did not induce neurotoxicity, even at maximum-tolerated doses of D-cycloserine.

**HIGHLIGHTS:**

- NRX-101, a fixed dose combination of D-cycloserine and lurasidone does not exhibit histologic neurotoxicity, even at maximum-tolerated doses

## Background

NRX-101 is a fixed dose combination of D-cycloserine and lurasidone that has been awarded Fast Track Designation and Breakthrough Therapy Designation by the US Food and Drug Administration (FDA) for the treatment of Severe Bipolar Depression with Acute Suicidal Ideation or Behavior. Preliminary evidence of efficacy for this unmet medical need was demonstrated in a phase 2 clinical trial (https://clinicaltrials.gov/ct2/show/NCT02974010) in which an 11 point improvement on the Montgomery Asberg Depression Rating Scale was demonstrated at day 14 (P=0.03). (Javitt 2018)

NRX-101 is now in phase 3 clinical trials under a Special Protocol Agreement awarded by the FDA. NRX-101 has additionally been awarded a Biomarker Letter of Support by the FDA (https://www.fda.gov/media/112706/download) based on demonstration of a reduction in glutamate in the prefrontal cortex on Magnetic Resonance Spectroscopy (MRS) and a statistically-significant correlation between this biomarker effect and reduced levels of depression. (Dong 2021) The purpose of this study is to demonstrate lack of neurotoxicity on histologic grounds as required by FDA in its recent guidance to industry related to development of novel antidepressants that target the N-methyl-d-aspartate receptor (NMDAR).

NMDAR antagonists are a new class of drugs that are rapidly gaining interest for the treatment of depression, PTSD, chronic pain, and other conditions. Unlike existing serotonin-targeted antidepressants, NMDAR-targeted antidepressants have rapid onset of action and may be suitable for use in patients with suicidal ideation. This class of drugs is now commonly referred to as Rapid Acting Antidepressants (RAADs) and are identified by FDA as having unique safety considerations based on potential neurotoxicity (FDA 2018). At the same time, the American Psychiatric Association has identified NMDAR-targeted RAADs as a promising new generation of drugs (Newport, et. al., 2015).

Despite their important clinical potential, NMDAR-targeted RAADs have been identified as having potential for neurotoxicity, a concern that has been echoed in an FDA safety communication (FDA 2017) and updated Guidance to Industry (FDA 2018) in which formal neurotoxicity studies are required prior to human trials of NMDAR-targeted antidepressants.

Excitatory neurotoxicity (Olney’s lesions) caused by ketamine was first reported in rodents in 1989 (Olney et al. 1989). Ketamine is one of the longest used and best-known NMDA antagonist drugs. Although it was shown to cause neuronal apoptotic lesions under experimental conditions, in usual clinical practice associated with single episodes of anesthesia it does not induce neurotoxicity. (Ding et al. 2016). MK-801 known as dizocilpine (INN) is an uncompetitive antagonist of NMDA receptor. It is known to produce Olney’s lesions in test rats. Because of its extreme propensity for neurotoxicity, it is commonly used as a positive control in drug safety studies.

Ketamine was serendipitously discovered to have potent and rapidly acting antidepressant effects (Berman et al., 2000, Aroni et al. 2009), thereby activating renewed scientific focus on NMDA antagonists as rapid-acting antidepressants. The antidepressant effect of DCS appears to be linked to its propensity to raise glutamate+glutamine (Glx) in the anterior cingulate cortex. Subnormal levels of Glx have been associated with depression in multiple case/control studies. (Luykx 2012). Cognitive disturbances are frequent in chronic abusers of ketamine, as well as frontal white matter abnormalities. Animal studies suggest that neuro-degeneration is a potential long-term risk of anesthetics in neonatal and young pediatric patients. (FDA Safety Communication 2017).

D-Cyloserine (DCS), an anti-infective drug used to treat tuberculosis was serendipitously discovered to have anti-depressant properties in 1959 (Crane 1959). However, because of its hallucinogenic potential it was contraindicated for use in treatment of depression. DCS has subsequently been identified as is a mixed agonist/antagonist of the NMDAR and appears to act via allosteric modification of the glycine-binding site. (Duncan et al. 2004). In 2009, Heresco-Levy and others reported that DCS when administered in conjunction with drugs that block the 5-HT2a receptor has an additive antidepressant effect with no observed hallucinogenic potential (Heresco-Levy, et. al 2009). In patients with bipolar depression, DCS has been shown to reduce glutamine levels in the in the brain in a manner comparable to ketamine and this reduction in glutamine correlates to decreased symptoms of depression, a finding also seen with ketamine. (Kantrowitz, et. al., 2021).

The phase 2 STABIL-B trial (https://clinicaltrials.gov/ct2/show/NCT02974010) was provided preliminary evidence of efficacy as required by the FDA for award of Breakthrough Therapy Designation. Subsequently, Chen (Chen et. al., 2019) reported that administration of DCS after ketamine infusion in patients with treatment resistant depression and suicidality was associated with lower scores on the Hamilton Depression Rating Scale item 3 (suicide), compared to patients treated with placebo (P=.01)

NRX-101 was developed as a novel oral antidepressant based on a fixed dose combination of DCS and lurasidone. The latter component is a D2/5HT_2A_ inhibitor that appears to block the hallucinatory side effects which might otherwise be seen with administration of DCS at high doses. While lurasidone’s precise mechanism of action is unknown its efficacy suggests involvement in the mediation of central dopamine type 2 and serotonin type 2 (5HT-2A) receptor antagonism.

This neurotoxicity study was designed as an IND-enabling study using a protocol agreed to with the FDA to demonstrate the neurological safety of increasing doses of DCS when combined with lurasidone by itself and following infusion of IV ketamine. Initial findings were presented at the National Biotechnology Conference of the American Association of Pharmaceutical Scientists (Kale 2018).

## 2. Materials and Methods

The study was performed under US and international Good Laboratory Practices (GLP) standards under the sponsorship of NeuroRx, Inc. (Wilmington, DE), according to a protocol reviewed and accepted in advance by the US Food and Drug Administration. Drug administration was performed by WuXi Apptec, (Shanghai, CN) under supervision of NeuroRx, Inc. Pathology specimens were prepared at Veterinary Pathology Services (Mason, OH) under our supervision (WJ) and examined by a single, experienced veterinary pathologist who was masked as to the treatment assignment of the animal subject. All experimental subjects were treated in accordance with the guidelines of the International Journal of Toxicology and their ethical treatment was overseen by the animal rights committee of WuXi Apptect, which operates in full accord with US and international law.

### 2.1 Subjects

A total of 106 female Sprague Dawley rats from Vital River Laboratory Animal Technology of China, 13 weeks of age and weighing 235.79-329.33 g, were housed up to 3 per cage and maintained on a 12 h light/dark cycle. Rats had unlimited access to water and food in their home cages. Animal subjects were acclimatized for 6 weeks prior to treatment. The study protocol was approved by the Animal Research Ethics Committee of China in accordance with China experimental animal administrative regulations. All efforts were made to minimize the number of animals used and to reduce their suffering. They were assigned to seven study groups, including Group 1 for vehicle control, Group 2 for positive control (MK-801), Groups 3 through 5 for ketamine+DCS/lurasidone (low, medium and high doses, respectively), Group 6 for ketamine alone, and Group 7 for DCS/lurasidone. Each group had 5,6,7 or 10 main study animals and 3 or 6 toxicokinetic (TK) animals (except the positive control groups).

### 2.2 Study Drug administration

Ketamine, DCS/lurasidone, and MK-801 were administered by IV infusion (40 minutes), oral gavage, and subcutaneous injection, respectively. Study drug levels and group assignments are identified in Table 1.

**TABLE 1.** Study drug dose, dosing volume and test article concentration for ketamine, DCS/lurasidone, and MK-801.

| Group | ketamine doses<sup>a</sup><br>(IV injection) |  |  | DCS/lurasidone doses<sup>a</sup><br>(Oral gavage) |  |  |
| --- | --- | --- | --- | --- | --- | --- |
|  | Dose<br>(mg/kg) | Volume<br>(mL/kg) | Conc.<br>(mg/mL) | Dose<br>(mg/kg) | Volume<br>(mL/kg) | Conc.<br>(mg/mL) |
| 1 | 0 | 0 | 0 | 0 | 0 | 0 |
| 2 | 0 | 0 | 0 | 0 | 0 | 0 |
| 3 | 35 | 0.7 | 50 | 500/35 | 10 | 50/3.5 |
| 4 | 35 | 0.7 | 50 | 1000/69 | 10 | 100/6.9 |
| 5 | 35 | 0.7 | 50 | 2000/139 | 10 | 200/13.9 |
| 6 | 35 | 0.7 | 50 | 0 | 0 | 0 |
| 7 | 0 | 0 | 0 | 2000/139 | 10 | 200/13.9 |

| Group | Vehicle<br>(IV injection) |  |  | MK-801<br>(SC injection) |  |  |
| --- | --- | --- | --- | --- | --- | --- |
|  | Dose<br>(mg/kg) | Volume<br>(mL/kg) | Conc.<br>(mg/mL) | Dose<br>(mg/kg) | Volume<br>(mL/kg) | Conc.<br>(mg/mL) |
| 1 | 0 | 10 | 0 | 0 | 0 | 0 |
| 8 | 0 | 0 | 0 | 10 | 2 | 5 |

Animals in Groups 3 through 5 were administered 12.5 mg/kg ketamine on Day 1, followed by 500/35 mg/kg, 1000/69 mg/kg, or 2000/139 mg/kg DCS/lurasidone on Day 2 (approximately 24 hours post ketamine infusion). Animals in Group 1 were administered the vehicle (50 mM Carbonate buffer pH 10) alone on Day 2 via oral gavage. Animals in Group 7 were administered 2000/139 mg/kg DCS/lurasidone only on Day 2. Animals in Group 2 and 8 were administered 1 and 5 mg/kgMK-801 on Day 1, respectively. Initially ketamine was dosed to 6 TK rats at 35 mg/kg via an IV bolus injection. Due to unscheduled deaths observed following this dose, the ketamine dose was reduced to 12.5 mg/kg and the dosing route was changed to IV infusion to mimic the clinical dosing route.

Animals were reassigned to replace those already dosed with 35 mg/kg ketamine. The day of ketamine administration was designated as Day 1 for all groups. All main study animals were sacrificed at 48 or 72 hours after ketamine administration. Main study animals were anesthetized by pentobarbital sodium and perfused with 10% neutral buffered formalin. The fixed brain and spleen tissues were collected and examined microscopically. Microscopic examinations included evaluations of H&E stained sections of brain and spleen and silver (Sevier-Munger modification of Bielschowsky’s method) and Fluoro-Jade B (FJB) stained sections of brain (Jordan and Hall 2007, Schmued and Hopkins 2000). Fourteen coronal planes of section of brain beginning with prefrontal cortex and extending through cerebellum and brain stem were evaluated by each stain. Blood samples were collected from TK animals to evaluate systemic exposure of ketamine and DCS/lurasidone.

### 2.3 Measurement of Study Drug levels

Toxokinetics for ketamine were assessed at baseline, 15 min, 1 hour, 4 hours, and 24 hours post IV infusion. Toxokinetics for NRX-101 were assessed at baseline, .5 hours, 1, 2, 4, 8, and 24 hours post oral gavage. At each time point, approximately 0.3 mL of blood was collected from animals from the jugular vein. Blood was collected into appropriately labeled tubes containing K2EDTA as the anticoagulant. The tubes were gently inverted several times to ensure mixing and immediately placed on ice. The quantification of ketamine, D-cycloserine, and lurasidone concentrations was conducted at the Testing Facility using validated liquid chromatographic triple quadrupole mass spectrometric (LC-MS/MS) methods (methods no. 409-0013-M for D-cycloserine and 409-0015-M for ketamine and lurasidone).

### 2.3 Regulatory Compliance

All portions of this study and histopathology evaluation were conducted in compliance with OECD Principles of Good Laboratory Practice, as revised in 1997 and adopted November 26th, 1997 by decision of OECD Council (C 97) 186/Final and US FDA Good Laboratory Practice Regulations 21 CFR 58, effective June 20, 1979, as amended 52 FR 33780, September 4, 1987, and subsequent amendments. National Institutes of Health guide for the care and use of Laboratory animals (NIH Publications No. 8023, revised 1978) were strictly observed in this study.

## 3. Results

Parameters evaluated during the study included mortality, clinical observations, body weights, food consumption, ophthalmology examinations, clinical pathology (hematology, coagulation, serum chemistry, and urinalysis), gross pathology, histopathology of brain and spleen, and toxicokinetics. The measured concentrations of DCS/lurasidone in the dosing formulations were 92%/94%, 83%/94% and 80%/72% of nominal values for the low-, mid-, and high-dose groups of DCS/lurasidone. The concentrations of DCS in the mid- and high-dose formulations and lurasidone in the high-dose formulation did not meet the acceptance criteria of 85% to 115%. These out-of-specification dose concentrations were not considered to have impacts on the overall objective achievement and conclusion of the study as the doses, when corrected by the actual concentrations, still gave a NOAEL of more than 10-fold safety margin for clinical trial.

After an IV infusion of ketamine on Day 1, Tmax values for Ketamine were observed at 0.3 hour post-dose. The Cmax values were observed between 871 and 1010 ng/mL, and the AUC_0-24h_ values were between 754 and 1160 h*ng/mL. After oral administration of DCS/lurasidone, Tmax values for DCS were observed at 0.5 and 1.0 hour post-dose. The Cmax values were observed between 378000 and 526000 ng/mL, and the AUC_0-24h_ values were between 938000 and 2950000 h*ng/mL. After oral administration of DCS/lurasidone, Tmax values for Lurasidone were observed at 2.0 to 24.0 hours post-dose. The Cmax values were observed between 301 and 636 ng/mL, and the AUC_0-24h_ values were between 2790 and7280 h*ng/mL.

### 3.1 Clinical Findings

Following oral dosing of DCS/lurasidone, test article-related clinical signs were observed with a dose-related incidence and included soiled coat at ≥500/35 mg/kg, ocular and/or nasal discharge at ≥1000/69 mg/kg, decreased activity and material around nose at 2000/139 mg/kg in animals dosed with or without ketamine, and prostration, atonia and closed eyes in animals dosed 2000/139 mg/kg DCS/lurasidone alone. Clinical signs observed in the positive control groups (MK-801) included prostration, atonia, decreased activity, abnormal gait, salivation ocular discharge and material around nose at ≥1 mg/kg.

Decreased body weight of 5 to 8% (relative to start weight) was noted following oral dosing of DCS/lurasidone at≥1000/69 mg/kg, which was accompanied by reduced food consumption of 31 to 77% (relative to concurrent control value) noted at ≥500/35 mg/kg (figs 1&2). Decreased body weight (10 to16%) (Figure 1) was noted in the MK-801 positive control group (Figure 1).

**Figure 1:**
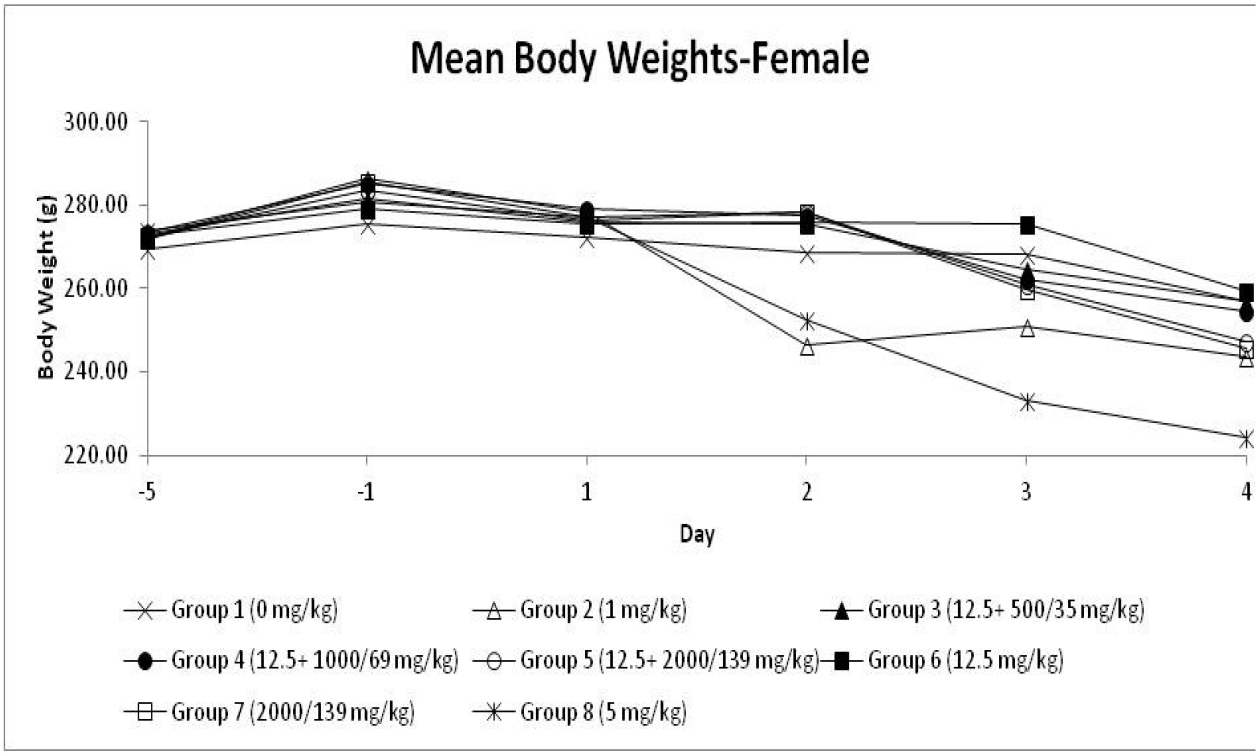
Mean Body Weight after administration of test article. A 10-16% reduction in body weight was noted in the MK=801 Positive Control Group (8).

**Figure 2:**
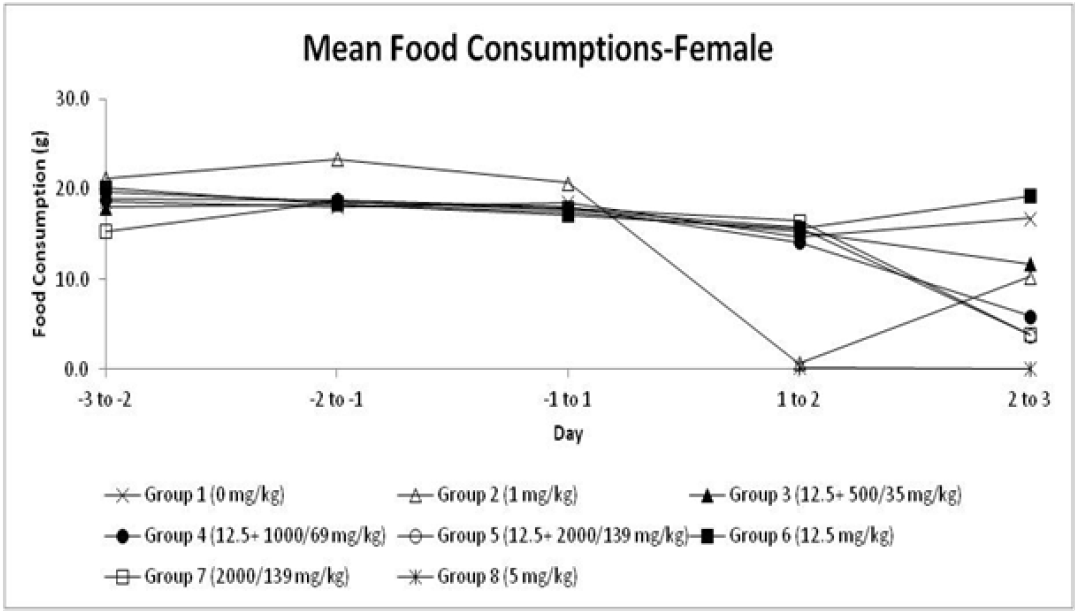
Mean Food Consumption after administration of test article. A moderate reduction in food consumption is noted at doses of D cycloserine in excess of 69 mg/kg (groups 4-7) with a >95% reduction in food consumption noted in the MK=801 Positive Control Group (group 8).

Decreased food consumption (∼50%) was seen with doses of D-cycloserine in excess of 69 mg/kg (groups 4-7). Complete cessation in food consumption, with lethargy was noted in the MK-801 positive control group 8 (Figure 2).

There were no test article-related morphological changes in ophthalmology examinations. Consistent with the clinical observations, red or brown ocular discharge and excessive lacrimation were principally noted in rats given 2000/139 mg/kg DCS/lurasidone with or without ketamine. Similar ocular findings were also observed in the positive control groups.

Test article-related hematology changes included increased neutrophils and decreased lymphocytes at ≥500/35 mg/kg DCS/lurasidone, and decreased reticulocyte count, increased eosinophils at 2000/139 mg/kg DCS/lurasidone with or without ketamine, and possibly increased platelets at ≥500/35 mg/kg DCS/lurasidone. Increased neutrophils and decreased lymphocytes were likely direct effects of the test article or related to a stress response. Decreased reticulocyte count was likely related to decreased food consumption. Increased neutrophils and decreased reticulocytes were also noted in the positive control groups. Test article-related coagulation changes were restricted to decreased prothrombin time noted at≥500/35 mg/kg DCS/lurasidone. Decreased prothrombin time and increased fibrinogen were noted in the 5 mg/kg MK-801 dose group.

Test article-related serum chemistry changes included increased serum total bilirubin at ≥1000/69 mg/kg DCS/lurasidone, increased serum urea in all DCS/lurasidone dose groups, increased serum triglyceride at 2000/139 mg/kg DCS/lurasidone with or without ketamine, and decreased serum alanine aminotransferase at≥1000/69 mg/kg DCS/lurasidone. Increased total bilirubin and triglyceride may suggest effects on the bile duct. Increased urea may suggest renal nephropathy. Decreased alanine aminotransferase was not considered of toxicological significance. Test article-related changes in urinalysis included increased urine protein, urine urobilinogen, and urine glucose at ≥1000/69 mg/kg DCS/lurasidone, which may suggest renal nephropathy. Increased urine protein and urine glucose were also noted in the 5 mg/kg MK-801 dose group.

### 3.2 Histopathology

For all subjects, the entire brain was removed, flushed so as to be clear of erythrocytes, fixed, embedded in paraffin, sectioned, stained with hematoxylin and eosin or Fluoro-jade, and examined microscopically (Schmued and Hopkins 2000, Switzer III et al., 2011). The brain tissue was examined for assessing potential Olney lesion according to procedures published in related literature (Olney JW.et al 1989a, Olney JW. et al 1991b). The brain was trimmed into 8 sections as described in the literature (Bolon et al 2013, Carson and Cappellano 2015). The blocks containing posterior cingulate and retrosplenial cortices were subjected to at least 3 non-serial sections for each level about 50-100 microns apart. At the discretion of the study pathologist, epifluorescent evaluation of stained microscopic sections was conducted to aid in interpretation of potential neurodegeneration, a procedure described by Jordan and Hall, 2007.

The sensitivity of the testing method was demonstrated by the presence of dose-responsive severity and distribution of neuronal degeneration/necrosis in Layers III and IV of the posterior cingulate and retrosplenial cortices and in other expected sub-anatomic brain locations of MK-801 positive control rats (figure 3). No microscopic abnormalities were observed in the brain of subjects given any dose of ketamine, DCS/lurasidone alone (figure 4), or ketamine followed by DCS/lurasidone (figure 5). The figures depict representative histology for the 2000 mg/Kg DCS with 139mg/Kg lurasidone (figure 4) and for 12.5 mg ketamine with 2000 mg/Kg DCS with 139mg/Kg lurasidone (figure 5). At 40x magnification using stains FJB and HE the combination of DCS/lurasidone with or without ketamine reveal normal neuronal structures and no evidence of neurotoxicity as shown by absence of degenerating neurons.

**Figure 3.**
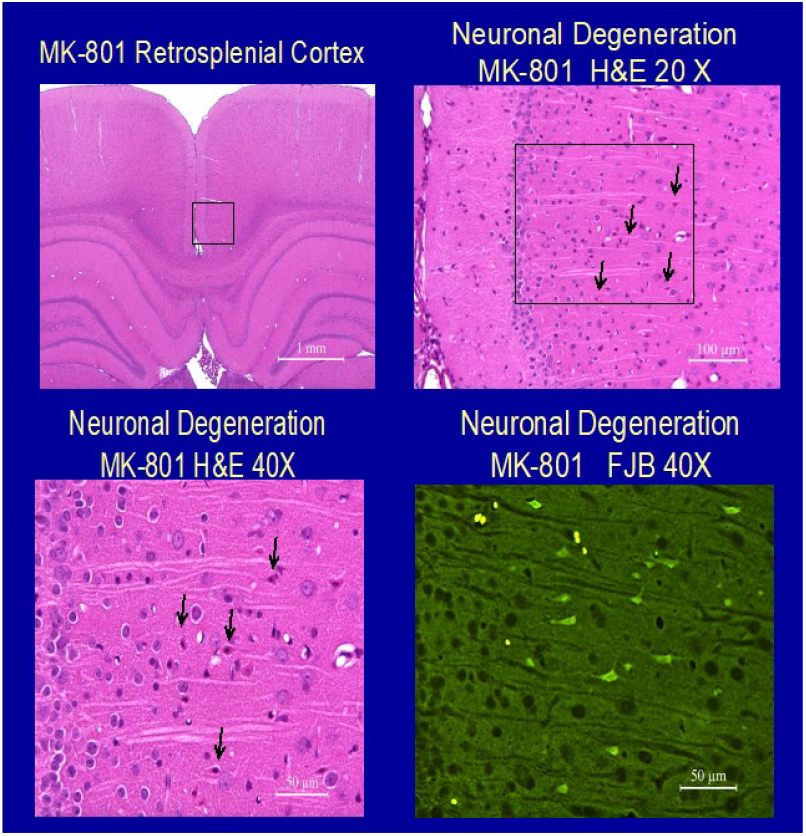
Representative histology in retrosplenial cortex for subjects treated with MK-801. Neuronal degeneration consistem with Olney’s lesions is seen (arrows)

**Figure 4:**
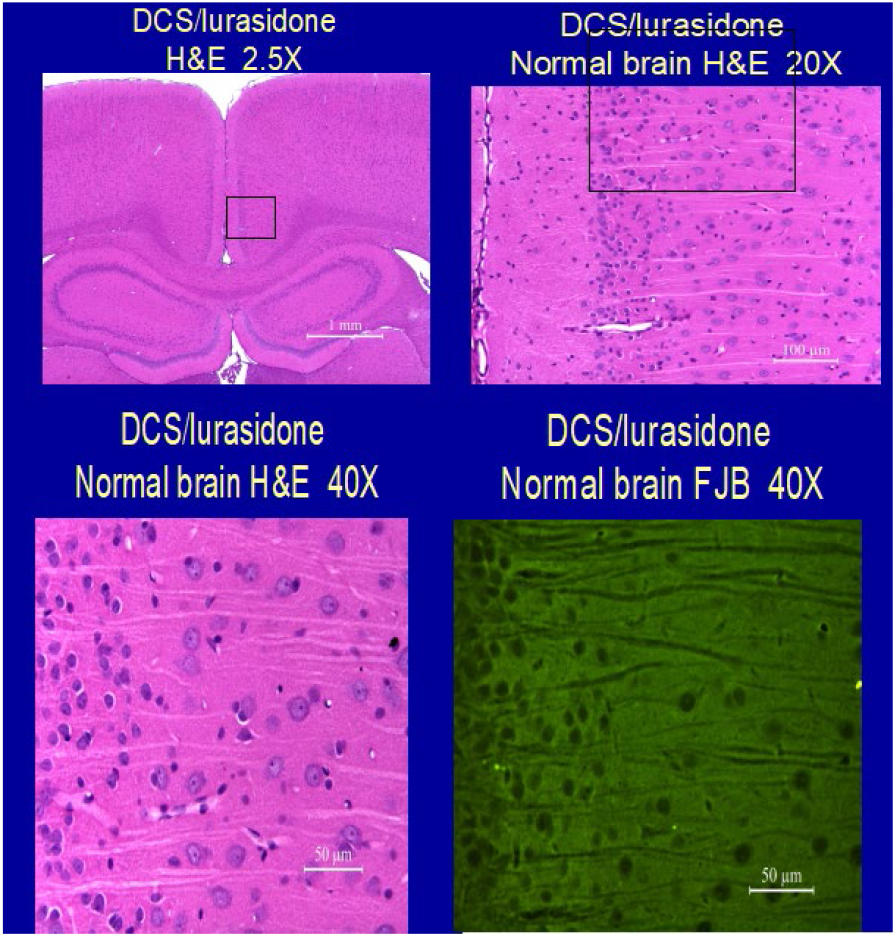
Representative histology in retrosplenial cortex for subjects treated with NRX-101. No evidence of Neuronal degeneration is seen.

**Figure 5:**
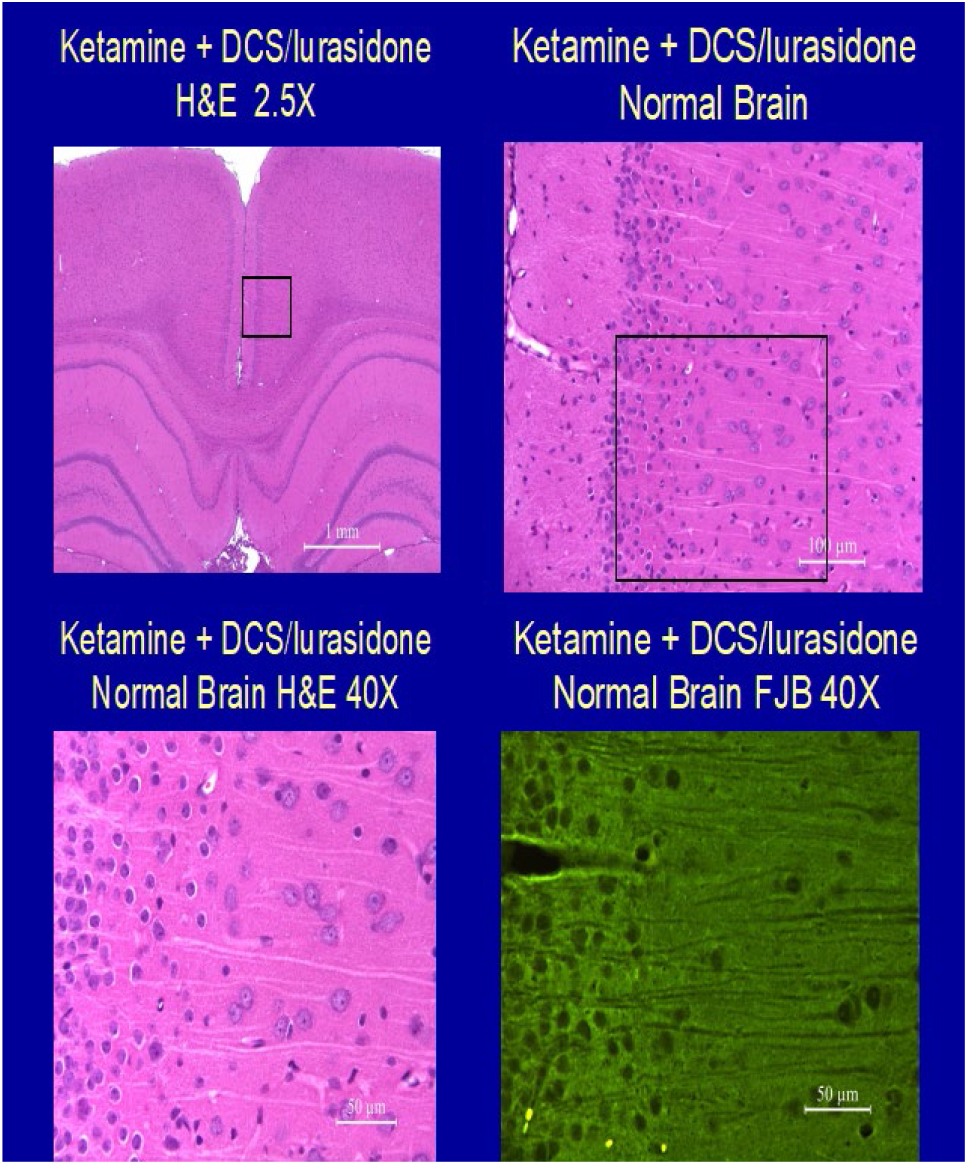
Representative histology in retrosplenial cortex for subjects treated with Ketamine followed by NRX-101. No evidence of Neuronal degeneration is seen.

The clinical signs observed in DCS/lurasidone-dosed animals, including decreased activity, prostration, atonia and closed eyes were considered extension of pharmacological effects of DCS and/or lurasidone. Other clinical signs observed, including ocular and/or nasal discharge, material around nose, and soiled coat were considered related to stress. Therefore these changes did not suggest additional neuronal toxicity of the treatment. Other findings including decreased body weight and food consumption, ocular changes, changes in clinical pathology were also not related to neuronal toxicity of the treatment. These results, together with the key study outcome that DCS/lurasidone treatment administered with or without ketamine did not induce microscopic abnormalities in the brain, suggest the intended treatment did not result in neuronal toxicity in this study. For all endpoints and parameters evaluated, changes seen in animals treated with 12.5 mg/kg ketamine +2000/139 mg/kg DCS/lurasidone were in general similar to those given 2000/139 mg/kg DCS/lurasidone alone, which suggests the combination treatment did not result in additional toxicity under the condition of the study.

## 4. Discussion

The data presented above support the FDA requirement to prove lack of histological neurotoxicity in association with novel NMDA-antagonist drugs that are proposed for the treatment of depression. The full data report associated with the above study was submitted to FDA and the agency issued a corresponding “Study may proceed” letter and subsequent “Special Protocol Agreement,” for phase 2b/3 clinical trials of NRX-101.

The findings demonstrate a substantial margin of safety as measured by histologic change associated in increasing doses of NRX-101. As doses increased from 500/35 to 2000/139 mg/kg, the systemic exposures (AUC_0-24h_ and/or Cmax) to DCS and lursidone increased dose-proportionally. At the same dose level of 2000/139 mg/kg DCS/lurasidone, the systemic exposures to DCS were comparable between animals dosed with or without ketamine; the systemic exposure to lurasidone was slightly lower in animals dosed without ketamine than in animals dosed with ketamine, which was likely due to a slightly extended Tmax. No obvious test-article related effects or changes were noted in animals given 12.5 mg/kg ketamine alone. In addition to the histologic safety, no clinical signs of neurotoxicity and no deaths of experimental subjects were observed.

The authors recognize that neurobehavioral testing and other techniques might in future be required to decisively prove the lack of neurotoxicity associated with NRX-101. However, the current FDA standard is based on histologic safety.

## 5. Conclusion

D-cycloserine/lurasidone combined, when administered once at doses up to 2000/139 mg/kg (actual dose 1600/100 mg/kg) to female rats via oral gavage with or without I.V. infusion of 12.5 mg/kg ketamine approximately 24 hours prior to the oral dosing, was tolerated and did not induce neurotoxicity. Under the conditions of the study, the No-Observed-Adverse-Effect Level (NOAEL) for neurotoxicity was considered to be 2000/139 mg/kg. The corresponding AUC_0-24h_ and C_max_ of DCS and lurasidone at the NOAEL were 2950000 h*ng/mL and 509000 ng/mL for DCS, and 7280 h*ng/mL and 636 ng/mL for lurasidone, respectively.

Based on these findings, the US FDA cleared NRX-101 for phase 2b3 trials in patients with bipolar depression and suicidal ideation.

